# Pathotyping of Newcastle disease virus: A novel single *BsaHI* digestion method of detection and differentiation of avirulent strains (lentogenic and mesogenic vaccine strains) from virulent virus

**DOI:** 10.1101/2021.07.16.452760

**Authors:** Perumal Arumugam Desingu, Shambhu Dayal Singh, Kuldeep Dhama, Obli Rajendran Vinodhkumar, K. Nagarajan, Rajendra Singh, Yashpal Singh Malik, Raj Kumar Singh

## Abstract

We provide a novel single restriction enzyme (RE) (*BsaHI*) digestion approach for detecting distinct pathotypes of the Newcastle disease virus (NDV). After scanning 4000 F gene nucleotide sequences in the NCBI database, a single RE (*BsaHI*) digesting site was discovered in the cleavage site. APMV-I “F gene” Class II specific primer-based reverse transcriptase PCR (RT-PCR) was utilized to amplify a 535 bp fragment, which was then digested with a single RE (*BsaHI*) for pathotyping avian NDV field isolates and pigeon paramyxovirus-1 isolates. The avirulent (lentogenic and mesogenic strains) produce 189 and 346 bp fragments, respectively, but the result in velogenic strains remains undigested with 535 bp fragments. In addition, 45 field NDV isolates and 8 vaccine strains were used to confirm the approach. The sequence-based analysis also agrees with the data obtained utilizing the single RE (*BsaHI*) digestion approach. The proposed technique had the potential to distinguish between avirulent and virulent strains in a short space of time, making it valuable in NDV surveillance and monitoring research.

## Introduction

Newcastle disease (ND) remains one of the most complex diseases to control in commercial and backyard poultry around the world. Newcastle disease virus (NDV) is a member of the genus *Avulavirus*, which belongs to the family *Paramyxoviridae*, under the order *Mononegavirales*. Avian paramyxoviruses (APMVs) are categorized into fifteen serotypes (APMV 1-15); with APMV-1 containing all economically relevant strains of NDV that occur naturally (1-4). APMV-1 is separated into two clades: class I and class II, with class II, subdivided into 21 genotypes (5-8). Class I isolates have a low virulence (lentogenic) and are mostly found in waterfowls (5, 7, 9, 10), while class II isolates have high virulence and are found in poultry, pets, and wild birds (5, 7, 9). The previous five panzootic outbreaks were caused by Class II viruses (7, 11-14). In addition, ND is divided into five pathotypes based on clinical signs and pathological lesions: 1) viscerotropic velogenic Newcastle disease (VVND) with digestive tract hemorrhagic lesions, 2) neurotropic velogenic Newcastle disease (NVND) with respiratory and neurological signs, and 3) mesogenic pathotype (a less pathogenic form of NVND), 4) Lentogenic pathotype with a mild or inapparent respiratory infection, and 5) Asymptomatic with no obvious disease. The intracerebral pathogenicity index (ICPI) in day-old chicks, as well as the demonstration of numerous basic amino acids at the fusion (F) protein cleavage site, are also required to report an epidemic caused by NDV (8) (OIE, Terrestrial manual, 2021). ICPI, on the other hand, is a time-consuming, expensive, arduous, and brutal procedure. Furthermore, due to the widespread use of the NDV vaccine strain or the presence of avirulent NDV strains in wild birds, the sequence analysis of the RT-PCR result is not suitable for all field instances.

Various quick techniques, such as RT-PCR followed by restriction enzyme (RE) analysis, were available with the development of molecular tools to discriminate avirulent and virulent ND virus isolates (15-18) and low virulent lentogenic field and vaccine strains were distinguished from mesogenic and velogenic field strains by RE digestion with B*gl*I. (19). However, most of them fail to differentiate mesogenic vaccine strains from velogenic strains. The *Bgl*I restriction site-based technique is one of the best techniques, whereas it had resulted in lentogenic sequences potentially misidentified as mesogenic and/or velogenic (20). The RT-PCR developed (21, 22) with specific reverse primers to differentiate NDV pathotype has a limited ability to amplify all virulent and avirulent strains of APMV-1 (including pigeon PMV-1). The probe hybridization (23, 24) and TaqMan™ fluorogenic probe hybridization (25) techniques to differentiate NDV pathotypes, however, these methods may not be befitting for routine laboratory diagnosis with limited facilities, and possibly increase the cost and time of the diagnosis.

In the present study, for the first time, we analyzed 4000 NDV sequences available in NCBI and identified a single RE site in the F gene cleavage site that can be used to differentiate avirulent (including lentogenic and mesogenic vaccine strains) NDV pathotypes from virulent strains. This appears to be the first report of RT-PCR-based single restriction enzyme digestion technique to differentiate avirulent, lentogenic, and mesogenic strains from virulent NDV strains for easy, economic, and specific diagnosis.

## Materials and Methods

### F gene cleavage site sequences based *in silico* determination of restriction enzyme

Newcastle disease virus F gene nucleotide and amino acid sequences were retrieved from GenBank NCBI (www.ncbi.nlm.nih.gov.in) and analysis of restriction enzyme sites at F gene cleavage site was done by using NEBcutter V2.0 (http://tools.neb.com/NEBcutter2/) and Webcutter 2.0 (http://rna.lundberg.gu.se/cutter2/) online software.

### Use of bioinformatics tools for identification of RE

The RE (*BsaHI*) was selected after sequence analysis of 4000 NDV F gene sequences. On alignment, all lentogenic strains of class II of APMV-1 contained *BsaHI* restriction site at the F0 cleavage region. This *BsaHI* site was missing at the F0 cleavage region from all mesogenic and velogenic strains selected for alignment. F gene amino acid ^119^GA^120^ adjacent to the cleavage site in mesogenic vaccine strains correspond to the *BsaHI* restriction site. However, no *BsaHI* cutting site is detected in velogenic strains.

### Validation of the technique

Newcastle disease virus isolates [45 field isolates from India and 8 vaccines (lentogenic and mesogenic) strains] obtained during 1989 to 2013 and maintained in the Virology Laboratory Repository, Avian Disease Section, Division of Pathology, Indian Veterinary Research Institute (IVRI), Izatnagar, India were used for this study. These isolates were reconstituted, and inoculums were prepared as explained by OIE, Terrestrial manual (2021), briefly 0.2 ml of inoculums was inoculated into 9-11-day-old SPF embryonated chicken eggs through allantoic route and the eggs were incubated at 37 °C till death or maximum period of 120 h, whichever is earlier. The embryos were candled every day and the dead embryos were chilled at 4 °C overnight and the collected allantoic fluids were tested for hemagglutination (HA) activity. After 120 h, all the remaining live embryos were collected and chilled at 4 °C for overnight. The allantoic fluids found negative for HA activity were further passage into at least one batch of eggs for confirmation. The harvested NDV was confirmed by the hemagglutination inhibition test and RT-PCR.

### RNA extraction and RT-PCR

Total RNA was extracted from allantoic fluid with TRIzol^R^ reagent (Cat. no. 15596-018, Invitrogen, USA) following the manufacturer"s instructions. The extracted RNA was used to synthesize cDNA by using random hexamer primer (Cat. no. 15596-018, MBI Fermentas, USA).

Reverse transcription (RT) for the first-strand synthesis was carried out by using RevertAid H minus (M-MuLV-RT) (MBI Fermentas, USA) in a standard 20 μl reaction mixture containing 5μl of total RNA, 1μl of random hexamer primer, 4.0 μl of 5X RT buffer, 2.0 μl of dNTP mix (10 mM each), 0.5 μl of RNAse inhibitor and 1.0 μl M-MLV RT enzyme (200 U/ μl). Reverse transcription was carried out at 25 ^0^C for 10 min; 42 ^0^C for 60 min and 70 ^0^C for 10 min. RT-PCR was done for all three pathotypes (lentogenic, mesogenic, and velogenic) with class II specific fusion (F) gene primers C2-F ATGGGCYCCAGACYCTTCTAC, C2-R CTGCCACTGCTAGTTGTGATAATCC (26). The RT-PCR was optimized in a standard 25 μl reaction mixture containing DNA/cDNA 3.0 μl, DreamTaq PCR Master mix 2X (Thermo scientific, India,) 12.5 μl, Forward primer (10 p mol/μl) 1.0 μl, Reverse primer (10 pmol/μl) 1.0 μl and Nucleus free water 7.5 μl. The cyclic conditions include an initial denaturation at 95°C for 4 min followed by 35 cycles of 94 °C for 40 sec, 52 °C for 50 sec, 72 °C for 50 sec and final extension at 72 °C for 5 min. The PCR products were analyzed by electrophoresis in 1.5% agarose gel stained with ethidium bromide (0.5µg/ml).

### Restriction endonuclease analysis

Class II specific F gene PCR products (535 bp) were analyzed by electrophoresis. PCR products were purified using the GeneJET PCR Purification kit (Thermo Scientific, USA) according to the manufacturer"s instructions. Purified PCR product was digested with *BsaHI* RE enzyme (NEB, UK), as per the manufacturer"s instructions. The reaction included NEB buffer 4 - 5.0 µl; BSA - 0.5 µl; PCR product - 10.0 µl (1 µg); *BsaHI* Enzyme - 1.0 µl (10 U); and nucleus free water - 33.5 µl. The mixture was incubated at 37°C for 2 to 3 h and the reaction was stopped by storing at -20 °C till further use.

### Sequencing and sequence analysis

The amplified Fgene PCR products were purified using GeneJET PCR purification kit (Thermo Scientific, USA) and sequenced through commercial sequencing service (BioServe, India). The F gene nucleotide sequences were translated by DNASTAR (version)/expasy translation tool (website) and the corresponding amino acid motif was identified.

## Results

### Identification of single restriction site (*BsaHI*) at F gene cleavage site

Of the 4000 NDV, F gene nucleotide, and amino acid sequences analyzed, 357 were class I specific sequences of APMV-1 and did not possess *BsaHI* restriction site within the F gene cleavage site. Thirteen amino acid motifs (^112^G/E/V/K-K/R/Q-Q-E/D/G-R/Q-L-I/V-G-A^120^) at the F gene cleavage site (**Supplementary Table 1**) were identified among these sequences. Motif patterns ^115^ERL^117, 115^DRL^117, 115^GRL^117^, and ^115^EQL^117^, were present in 91.32 %, 4.76 %, 3.64 % and 0.28 % of the sequences analyzed, respectively. All the patterns had "L" at 117 positions in the F gene cleavage site. The 858 Class II avirulent NDV sequences when analyzed showed 13 amino acid motifs (^112^G/E/R-K/R/T-Q-G/K/A/R-R-L-I/L-G-A/F^120^) with the patterns ^115^GRL^117, 115^KRL^117, 115^ARL^117,^ and ^115^RRL^117^ present in 98.94 %,0.82 %, and 0.12 % of the sequences, respectively. These 13 different amino acid motif patterns have 42 (**Supplementary Table 2 & 3**) different nucleic acid patterns and have at least one *BsaHI* digestion site. The nucleotide sequence - GG(A/G)CG(T/C)CT(T/G/C), corresponding to the amino acid pattern ^115^G/R-R-L^117^ at F gene cleavage site in class II specific avirulent and lentogenic vaccine strain’s sequence of APMV-1 has a *BsaH1* digestion site (GGCGCC). The motif ^115^K/A-R-L^117^ present in seven sequences (in less than 0.94 % of class II avirulent sequences) from India did not have a *BsaHI* digestion site at the F gene cleavage site, but ^119^GA^120^ pattern was noticed adjacent to the F gene cleavage site. Out of all the class II specific avirulent NDV, 54.19% have *BsaHI* digestion sites at both in and adjacent to the F gene cleavage site and the remaining 45.81% sequences showed *BsaHI* digestion site at only the F gene cleavage site. The analysis of 2584 sequences of Class II virulent NDV strains showed 19 different amino acid motifs (^112^R/K/G-R/R-Q/K/R/-K/R/G-R-F-I/V/L-G/S-A^120^) with the patterns ^115^KRF^117, 115^RRF^117,^ and ^115^GRF^117^ in 89.97 %, 9.99 %, and 0.04 % of the sequences analyzed, respectively. The 19 different amino acid motif patterns have 134 different nucleic acid patterns (**Supplementary table 4 & 5**) with no *BsaH1* digestion site.

It is of note that the mesogenic strains had the same motif pattern at the F gene cleavage site as the pathogenic/velogenic Class-II APMV-I. The mesogenic and velogenic strains also had the same amino acid motif ^119^GA^120^ adjacent to the cleavage site, but in mesogenic strains, the nucleotide sequence at this region corresponds to the *BsaH1* digestion site (GGCGCC). In velogenic strains, the nucleotide sequence at this region was found to be GG(C/T) GC(C/T) (**Supplementary Table 6 & 7**) with at least had one T in the degenerate nucleotide position.

### Relationship between sequence analysis and ICPI pathotyping

The amino acid motif at the F gene cleavage site, nucleotide pattern at F gene cleavage, and ICPI were analyzed for 212 sequences/strains. The *BsaH1* digestion of these sequences showed that, in 200 sequences, the ICPI matched with the availability of RE digestion site; but 12 (12/212) sequences did not match (**Supplementary Table 8**).

### Relationship between sequence analyses with MDT Pathotyping

The 82 NDV isolates/ strains sequences with MDT, for which amino acid motif at F gene cleavage site, nucleotide pattern at F gene cleavage site were accessible, when subjected to *BsaH1* analysis accurately matched (**Supplementary Table 9**).

### Vaccine strains

All NDV vaccine strains except Mukteswar and H vaccine strain’s sequences showed *BsaH1* digestion site at F gene cleavage (**Supplementary Table 10**).

### Validation of *in silico* analysis in laboratory condition

In this study, to validate the *in-silico* analysis results, 45 field isolates from India and commonly used eight vaccines (lentogenic and mesogenic) strains were used, where all the isolates and vaccine strains were found positive for 535 bp Class II specific APMV-1 RT-PCR. The avirulent, lentogenic, and mesogenic strains, excluding Mukteshwar, provided digested fragments when the Class II specific RT-PCR products of 535 bp were exposed to *BsaHI* digestion, however, the virulent NDV RT-PCR product remained undigested by this enzyme (**Figure 1 and Table 1**)

**Figure 1:**
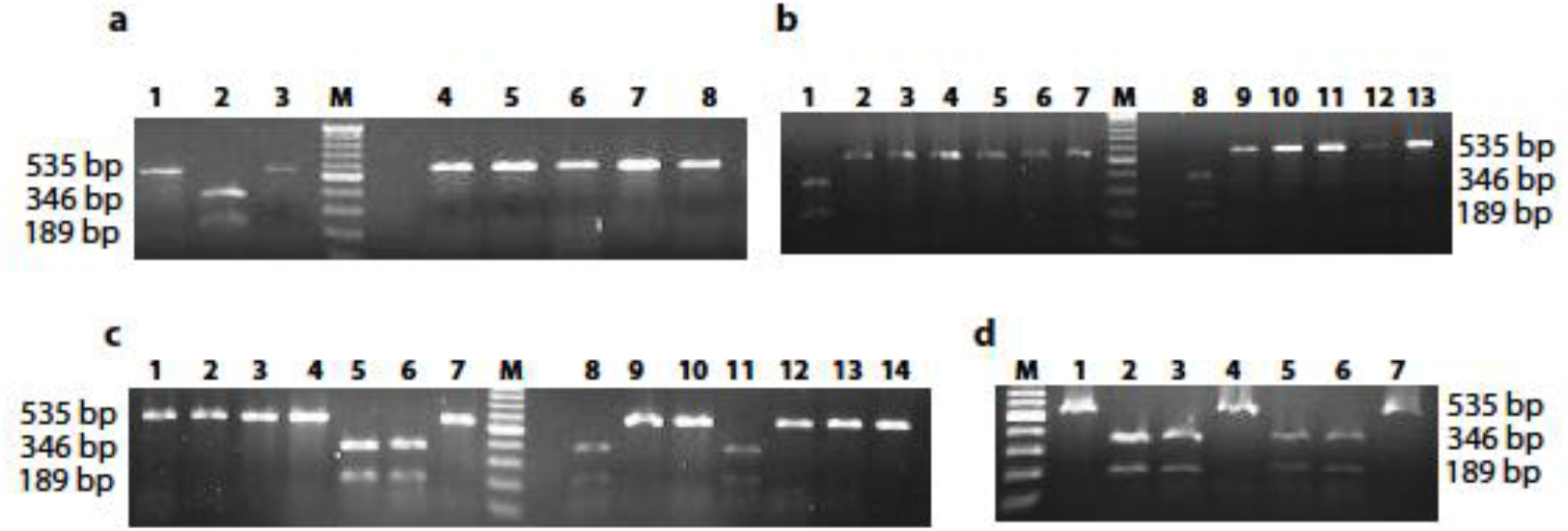
NDV F gene digestion products visualized by agarose gel electrophoresis. Totally 43 isolates were tested in this study including vaccine strains of NDV and other field isolates. All avirulent and mesogenic vaccine strains (except Mukteswar strain) of NDV were digested by *Bsa H1* RE enzyme and same time virulent strains of NDV were not digested, the agarose gel electrophoresis results are presented in **fig.1a-d. a**) M-Ladder; Lane1- 1/chicken/IVRI/13; Lane2- Crane/ADS/IVRI/13; Lane3- Chicken-India-UP-IVRI-0011-2010; Lane4- Emu-India-AP-IVRI- 0007-2010; Lane5- Pi04/AD/94; Lane6- Chicken-India-TN-IVRI-0005-2010; Lane7- IVRI-0010; and Lane8- 129/FAIZABAD/AD-IVRI/92. **b**) M- Ladder; Lane1- Hitchner B1; Lane2- 127/FAIZABAD/AD-IVRI/92; Lane3- IVRI/91; Lane4- IVRI/92; Lane5-123/KATHUA/AD- IVRI/92; Lane6- 75/RAMPUR/AD-IVRI/89; Lane7- 108/BAREILLY/AD-IVRI/91; Lane8- 3/chicken/IVRI/13; Lane9- Pi01/AD/91; Lane10- Pi02/AD/91; Lane11- Pi05/AD/97; Lane12- Pi06/AD/01; and Lane13- PiAD388/Quail/India. **c**) M-Ladder; Lane1- NDV/Peafowl/Haryana/India/IVRI-037/12; Lane2- NDV/Peafowl/UP/IVRI-024/12; Lane3- NDV/Peafowl/Delhi/IVRI-0022/12; Lane4- Mukteshwar; Lane5- Lasota; Lane6- R2B; Lane7- PiAD33/Guinea fowl/India; Lane8- Komarov; Lane9- PiAD18/Guinea fowl/India; Lane10- PiAD55/Pigeon/India; Lane11- F strain; Lane12- Chicken-India-TN-IVRI-0006-2010; Lane13- Chicken-India-TN-IVRI-0001-2010; and Lane14- Chicken-India-TN-IVRI-0003-2010. **d**) M- ladder; Lane1- PiADPrd/Pigeon/India; Lane2- 2/chicken/IVRI/13; Lane3- Roakin; Lane4- Chicken-India-TN-IVRI-0002-2010; Lane5- Beaudette C; Lane6- CDF 66; and Lane7- Chicken- India-KA-IVRI-0008-2010.

**Table 1:**
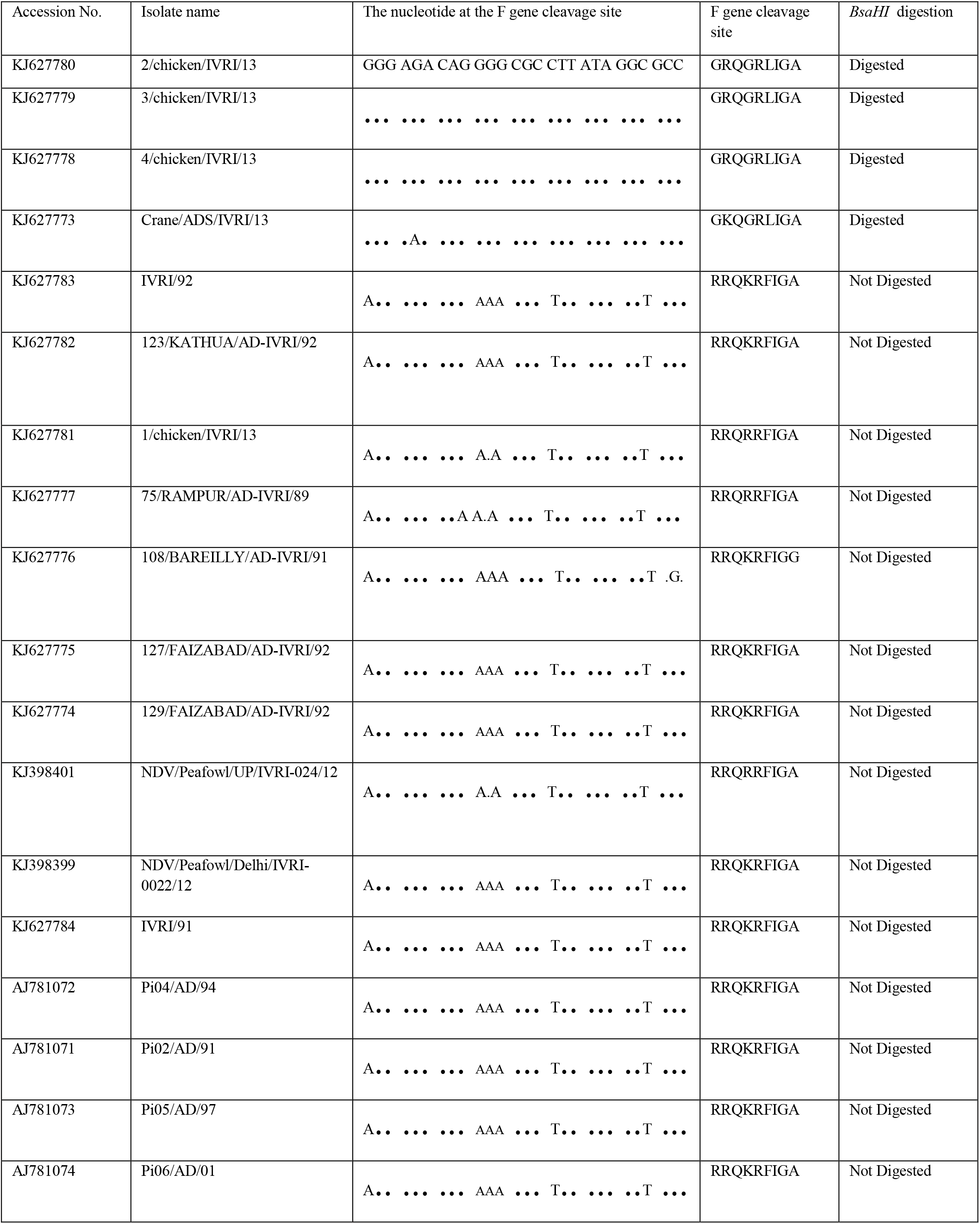
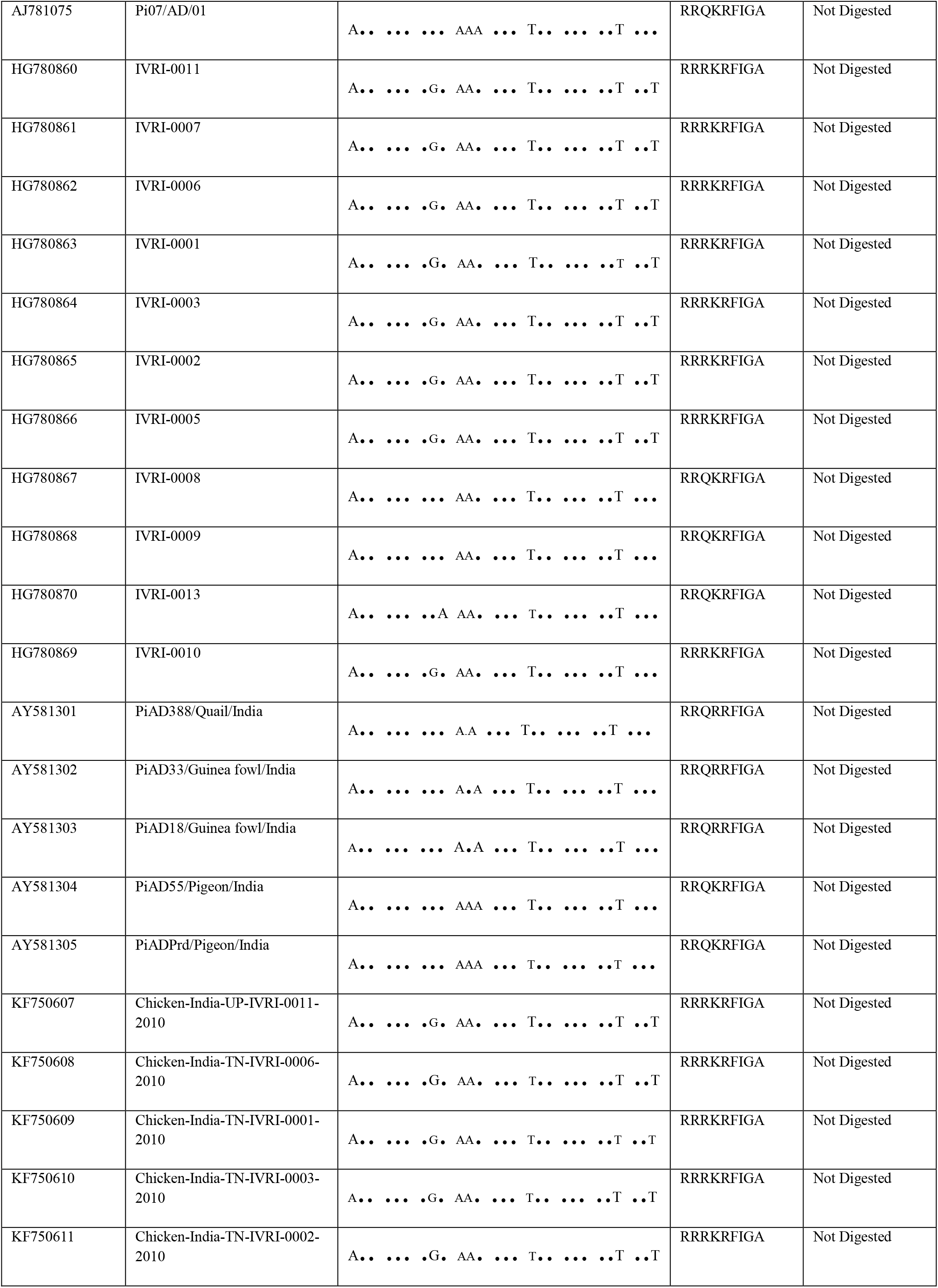
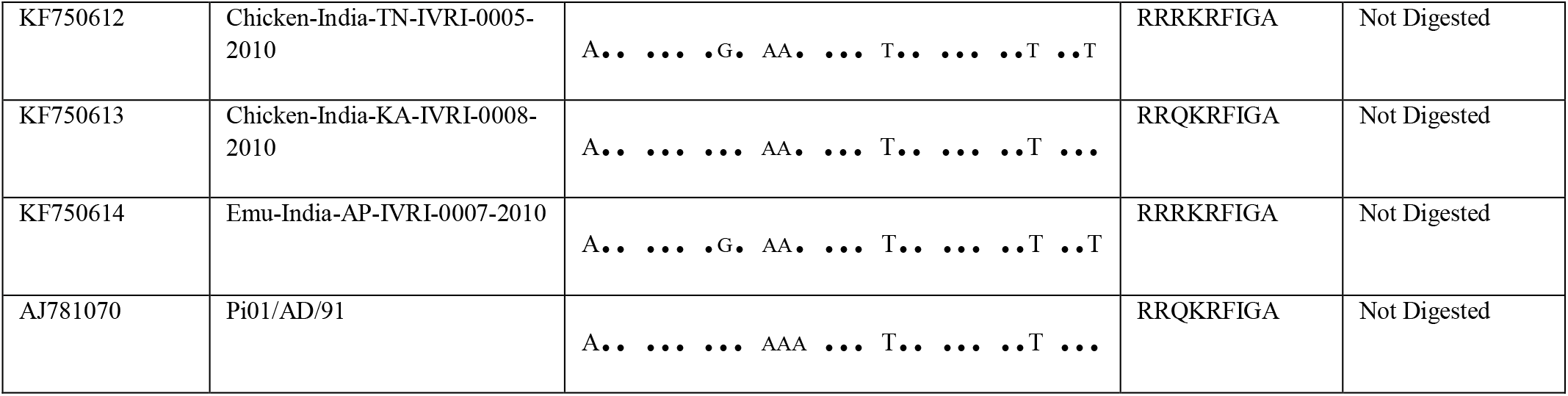
Comparison of F gene cleavage site amino acid motif and *BsaHI* digestion for detection of NDV pathotype based on 45 NDV field isolates used in this study.

The F gene PCR product of all the 44 field isolates was sequenced and submitted to the GenBank (Table 1). These sequences were aligned with 8 vaccine strain sequences to identify the F gene cleavage site, amino acid motif across sequences. In comparison, *BsaHI* digestion patterns across sequences with the F gene cleavage site, a 100% correlation was observed between F gene cleavage site amino acid motif-based pathotyping and *BsaHI* digestion pattern-based pathotyping.

## Discussion

The necessity of precise and timely diagnosis of NDV outbreaks cannot be overstated since the disease is economically important and transboundary. Currently recommended methods of pathotyping of NDV include the determination of intracerebral pathogenicity index (ICPI) and demonstration of the amino acid motif at the F gene cleavage site (OIE, Terrestrial manual, 2021). ICPI is time-consuming, labor-intensive, inhumane, and has limited use in surveillance programs. The routine PCR-based diagnosis of NDV is non-confirmatory and gives false-positive results as many avirulent strains of the wild population and the lentogenic vaccine strains used were re-isolated for several weeks post-administration of vaccine which may give false-positive results (19, 27). For pathotypic characterization of NDV isolates, the nucleotide variation around the F gene cleavage site has been extensively exploited and considered as the primary molecular determinant for NDV virulence (8, 25). However, demonstration of specific amino acid motif pattern in F gene cleavage site is costly, time-consuming, need sophisticated instruments, and has restricted use for field cases and surveillance. The available diagnostic methods warrant the need for accurate, rapid, simple, and economical tests for detection and differentiation of NDV pathotypes, which is important for surveillance and control of the disease during outbreaks. In the present study on *in-silico* analysis of 4000 sequences, the RE *BsaHI* was identified as the RE enzyme with a cutting site either within or adjacent to the F0 cleavage site to differentiate various pathotypes. The utility of the enzyme for pathotyping was validated on 45 field isolates and 8 vaccine strains.

The class I APMV-1 includes isolates of low virulent strains, and class II APMV-1 includes highly virulent, all vaccines and reasonable numbers of avirulent strains (26). Because almost all class I NDV isolates, except one, IECK90187 (10, 26) are lentogenic strains, our work relied on Class II specific detection. The most often utilized lentogenic and mesogenic vaccine strains are from APMV-I class II. From 1920 until the present, Class II APMV-I viruses were responsible for all five panzootic of NDV (12-14). The selection of the *BsaHI* restriction enzyme is based on the nucleotide sequence of avirulent, lentogenic, and mesogenic vaccine strains. The velogenic class-II velogenic strains did not have a *BsaHI* digestion site either within or adjacent to the F0 cleavage site. The APMV-I class II avirulent, lentogenic, and mesogenic vaccine strains that have the *BsaHI* cutting site were digested within or adjacent to the F0 cleavage site. However, the Mukteswar and H strains as vaccine strains belong to Class II genotype III of APMV-1. These strains do not contain the *BsaHI* RE site and have a GGTGCC nucleotide sequence at the site corresponding to the *BsaHI* cutting site within the F cleavage site. This indicates that these strains are velogenic strains adopted to be mesogenic vaccine strains by serial egg passages (28).

On analyzing the sequences vis-a-vis the available data on NDV pathotypes based on ICPI and MDT and comparing the amino acid motif at the cleavage site and the nucleotide pattern at the F gene cleavage site, 12 (12/278) cases were found to be mismatching between them. In any mismatching cases between amino acid motif at cleavage site and pathogenicity index (ICPI), the ICPI stands best to declare about the pathogenicity (OIE, Terrestrial manual, 2021). Though the amino acid motif at F gene cleavage site ^113^RXR/KRF^117^ is considered specific for virulent NDV pathotypes, the motif pattern of ^112^R-R-K-K-R-F^117^ in pigeon variant PMV-1 isolates have been correlated to both high and low virulence isolates/strains in ICPI tests conducted in chickens (29). The virulent motif ^112^G-R-Q-K-R-F^117^ of PPMV-1 isolates exhibited low virulence in chickens (30, 31).

There is a high similarity between the virulent, vaccine, and avirulent NDV strains, which hinders the diagnosis of NDV. Recently, Liu et al. (26) demonstrated the specificity of class II specific primers in sixty-seven field isolates tested. In our study, all the isolates identified as Class II APMV-I were confirmed as class II of NDV using class II specific RT-PCR (26). On sequencing 535 bp sequence amplified by RT-PCR in the 45 field isolates and comparing the F cleavage site motif pattern with the available vaccine strain sequences considered for validation, the results corroborated the findings from restriction enzyme analysis except for the Mukteswar vaccine strain. This novel method of RE (*BsaHI*) analysis is useful for both APMV-1 and PPMV-1. Furthermore, standard sequencing is unable to distinguish between mesogenic and velogenic strains at the cleavage location, but in this new procedure, one could differentiate mesogenic and velogenic strains. In a prior study, we used degenerate primers and nested RT-PCR to distinguish mesogenic and avirulent NDV strains (22).

## Conclusion

NDV pathotyping using the RT-PCR coupled RE digestion approach is fast, cheap, and ethical, allowing for large-scale surveillance and detection of virulent and avirulent as well as vaccine strains of NDV.

## Acknowledgments

The authors are highly thankful to the Indian Veterinary Research Institute, Izatnagar, and Indian Council of Agriculture Research, New Delhi for providing the necessary facilities to carry out this research work. P.A.D is a DST-INSPIRE faculty is supported by research funding from the Government of India (DST/INSPIRE/04/2016/001067), and the Science and Engineering Research Board, Department of Science and Technology, Government of India (CRG/2018/002192).

## Un-cropped original images

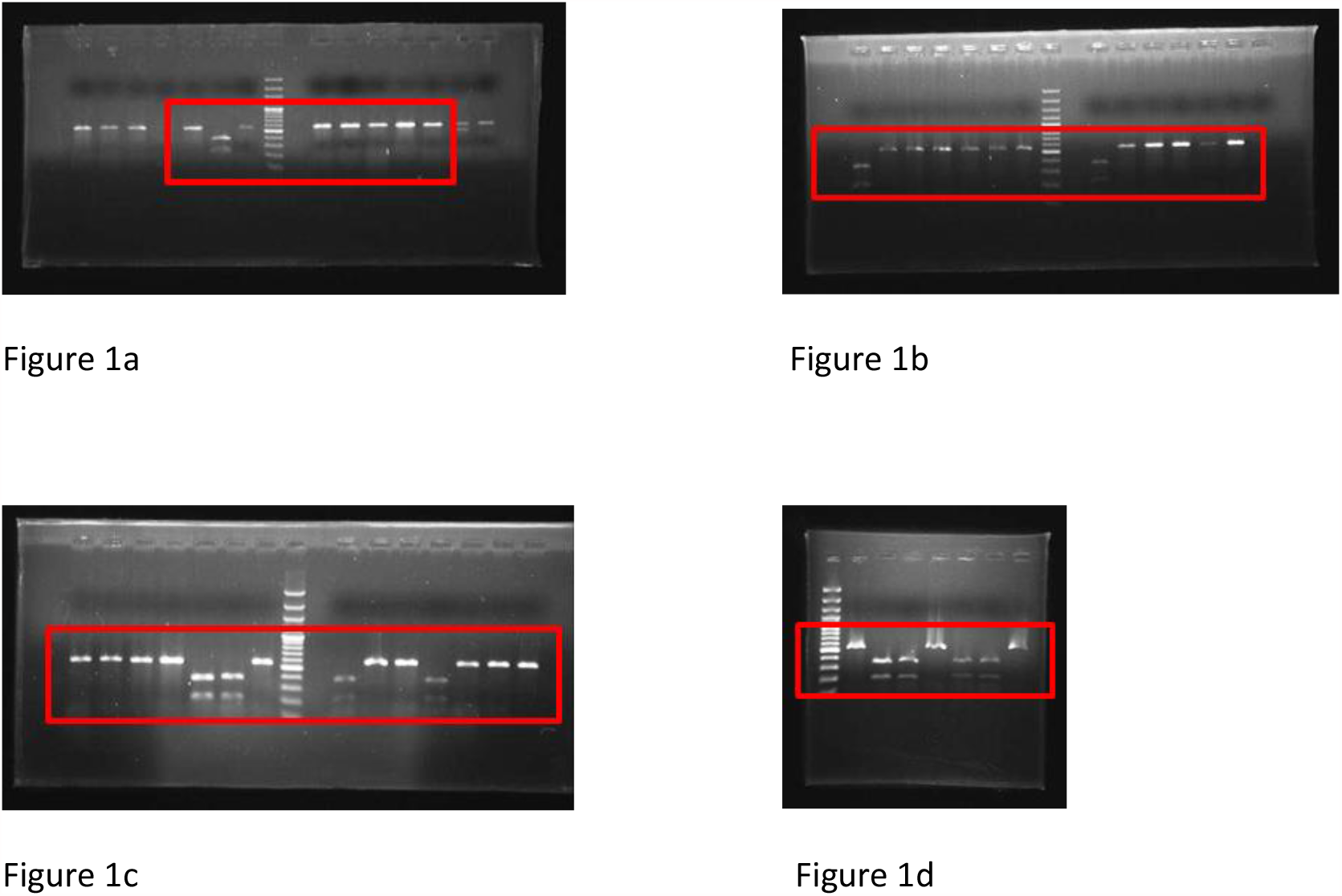

## References

1. Lee, H.J., Kim, J.Y., Lee, Y.J., Lee, E.K., Song, B.M., Lee, H.S., and Choi, K.S. 2017. A Novel Avian Paramyxovirus (Putative Serotype 15) Isolated from Wild Birds. Front Microbiol 8:786.

2. Fereidouni, S., Jenckel, M., Seidalina, A., Karamendin, K., Beer, M., Starick, E., Asanova, S., Kasymbekov, E., Sayatov, M., and Kydyrmanov, A. 2018. Next-generation sequencing of five new avian paramyxoviruses 8 isolates from Kazakhstan indicates a low genetic evolution rate over four decades. Arch Virol 163:331–336.

3. Nayak, B., Dias, F.M., Kumar, S., Paldurai, A., Collins, P.L., and Samal, S.K. 2012. Avian paramyxovirus serotypes 2-9 (APMV-2-9) vary in the ability to induce protective immunity in chickens against challenge with virulent Newcastle disease virus (APMV-1). Vaccine 30:2220–2227.

4. Terregino, C., Aldous, E.W., Heidari, A., Fuller, C.M., De Nardi, R., Manvell, R.J., Beato, M.S., Shell, W.M., Monne, I., Brown, I.H., et al. 2013. Antigenic and genetic analyses of isolate APMV/wigeon/Italy/3920-1/2005 indicate that it represents a new avian paramyxovirus (APMV-12). Arch Virol 158:2233–2243.

5. Liu, Y.P., Chang, C.Y., Lee, F., Chiou, C.J., and Tsai, H.J. 2020. Phylogenetic analysis of avian paramyxoviruses 1 isolated in Taiwan from 2010 to 2018 and evidence for their intercontinental dispersal by migratory birds. J Vet Med Sci 82:1366–1375.

6. Hicks, J.T., Dimitrov, K.M., Afonso, C.L., Ramey, A.M., and Bahl, J. 2019. Global phylodynamic analysis of avian paramyxovirus-1 provides evidence of inter-host transmission and intercontinental spatial diffusion. BMC Evol Biol 19:108.

7. Xiang, B., Chen, L., Cai, J., Liang, J., Lin, Q., Xu, C., Ding, C., Liao, M., and Ren, T. 2020. Insights into Genomic Epidemiology, Evolution, and Transmission Dynamics of Genotype VII of Class II Newcastle Disease Virus in China. Pathogens 9.

8. Desingu, P.A., Singh, S.D., Dhama, K., Vinodhkumar, O.R., Barathidasan, R., Malik, Y.S., Singh, R., and Singh, R.K. 2016. Molecular characterization, isolation, pathology and pathotyping of peafowl (Pavo cristatus) origin Newcastle disease virus isolates recovered from disease outbreaks in three states of India. Avian Pathol 45:674–682.

9. Kim, L.M., King, D.J., Curry, P.E., Suarez, D.L., Swayne, D.E., Stallknecht, D.E., Slemons, R.D., Pedersen, J.C., Senne, D.A., Winker, K., et al. 2007. Phylogenetic diversity among low-virulence newcastle disease viruses from waterfowl and shorebirds and comparison of genotype distributions to those of poultry-origin isolates. J Virol 81:12641–12653.

10. Alexander, D.J., Campbell, G., Manvell, R.J., Collins, M.S., Parsons, G., and McNulty, M.S. 1992. Characterisation of an antigenically unusual virus responsible for two outbreaks of Newcastle disease in the Republic of Ireland in 1990. Vet Rec 130:65–68.

11. Miller, P.J., Haddas, R., Simanov, L., Lublin, A., Rehmani, S.F., Wajid, A., Bibi, T., Khan, T.A., Yaqub, T., Setiyaningsih, S., et al. 2015. Identification of new sub-genotypes of virulent Newcastle disease virus with potential panzootic features. Infect Genet Evol 29:216–229.

12. Liu, H., Wang, Z., Wu, Y., Zheng, D., Sun, C., Bi, D., Zuo, Y., and Xu, T. 2007. Molecular epidemiological analysis of Newcastle disease virus isolated in China in 2005. J Virol Methods 140:206–211.

13. Miller, P.J., Decanini, E.L., and Afonso, C.L. 2010. Newcastle disease: evolution of genotypes and the related diagnostic challenges. Infect Genet Evol 10:26–35.

14. Miller, P.J., Kim, L.M., Ip, H.S., and Afonso, C.L. 2009. Evolutionary dynamics of Newcastle disease virus. Virology 391:64–72.

15. Ballagi-Pordany, A., Wehmann, E., Herczeg, J., Belak, S., and Lomniczi, B. 1996. Identification and grouping of Newcastle disease virus strains by restriction site analysis of a region from the F gene. Arch Virol 141:243–261.

16. Kou, Y.T., Chueh, L.L., and Wang, C.H. 1999. Restriction fragment length polymorphism analysis of the F gene of Newcastle disease viruses isolated from chickens and an owl in Taiwan. J Vet Med Sci 61:1191–1195.

17. Nanthakumar, T., Kataria, R.S., Tiwari, A.K., Butchaiah, G., and Kataria, J.M. 2000. Pathotyping of Newcastle disease viruses by RT-PCR and restriction enzyme analysis. Vet Res Commun 24:275–286.

18. Wehmann, E., Herczeg, J., Ballagi-Pordany, A., and Lomniczi, B. 1997. Rapid identification of Newcastle disease virus vaccine strains LaSota and B-1 by restriction site analysis of their matrix gene. Vaccine 15:1430–1433.

19. Creelan, J.L., Graham, D.A., and McCullough, S.J. 2002. Detection and differentiation of pathogenicity of avian paramyxovirus serotype 1 from field cases using one-step reverse transcriptase-polymerase chain reaction. Avian Pathol 31:493–499.

20. Cattoli, G., Susta, L., Terregino, C., and Brown, C. 2011. Newcastle disease: a review of field recognition and current methods of laboratory detection. J Vet Diagn Invest 23:637–656.

21. Kant, A., Koch, G., Van Roozelaar, D.J., Balk, F., and Huurne, A.T. 1997. Differentiation of virulent and non-virulent strains of Newcastle disease virus within 24 hours by polymerase chain reaction. Avian Pathol 26:837–849.

22. Desingu, P.A., Singh, S.D., Dhama, K., Kumar, O.R., Singh, R., and Singh, R.K. 2015. A rapid method of accurate detection and differentiation of Newcastle disease virus pathotypes by demonstrating multiple bands in degenerate primer based nested RT-PCR. J Virol Methods 212:47–52.

23. Jarecki-Black, J.C., Bennett, J.D., and Palmieri, S. 1992. A novel oligonucleotide probe for the detection of Newcastle disease virus. Avian Dis 36:134–138.

24. Raghavan, V.S., Kumanan, K., Thirumurugan, G., and Nachimuthu, K. 1998. Comparison of various diagnostic methods in characterizing Newcastle disease virus isolates from Desi chickens. Trop Anim Health Prod 30:287–293.

25. Aldous, E.W., and Alexander, D.J. 2001. Detection and differentiation of Newcastle disease virus (avian paramyxovirus type 1). Avian Pathol 30:117–128.

26. Liu, H., Zhao, Y., Zheng, D., Lv, Y., Zhang, W., Xu, T., Li, J., and Wang, Z. 2011. Multiplex RT-PCR for rapid detection and differentiation of class I and class II Newcastle disease viruses. J Virol Methods 171:149–155.

27. van Eck, J.H., van Wiltenburg, N., and Jaspers, D. 1991. An Ulster 2C strain-derived Newcastle disease vaccine: efficacy and excretion in maternally immune chickens. Avian Pathol 20:481–495.

28. Czegledi, A., Wehmann, E., and Lomniczi, B. 2003. On the origins and relationships of Newcastle disease virus vaccine strains Hertfordshire and Mukteswar, and virulent strain Herts’33. Avian Pathol 32:271–276.

29. Werner, O., Romer-Oberdorfer, A., Kollner, B., Manvell, R.J., and Alexander, D.J. 1999. Characterization of avian paramyxovirus type 1 strains isolated in Germany during 1992 to 1996. Avian Pathol 28:79–88.

30. Collins, M.S., Strong, I., and Alexander, D.J. 1996. Pathogenicity and phylogenetic evaluation of the variant Newcastle disease viruses termed “pigeon PMV-1 viruses” based on the nucleotide sequence of the fusion protein gene. Arch Virol 141:635–647.

31. Zanetti, F., Mattiello, R., Garbino, C., Kaloghlian, A., Terrera, M.V., Boviez, J., Palma, E., Carrillo, E., and Berinstein, A. 2001. Biological and molecular characterization of a pigeon paramyxovirus type-1 isolate found in Argentina. Avian Dis 45:567–571.

